# Poorer physical function is associated with elevated spatial entropy in the aging brain network landscape

**DOI:** 10.64898/2025.12.09.693258

**Authors:** Clayton C. McIntyre, Shannon M. O’Donnell, Robert G. Lyday, Jonathan H. Burdette, Steven R. Cummings, Stephen B. Kritchevsky, Paul J. Laurienti

## Abstract

Life is a constant struggle against disorder. As we age, our ability to maintain internal order declines. In the healthy human brain, order is observable in the form of functionally segregated brain network communities that exhibit spatial consistency. These communities associate with distinct cognitive and physical functions. When mapped into the brain, they form a functional “landscape”. We assessed the spatial disorder of these landscapes in older adults with a wide range of mobility using a modified version of Shannon entropy. We found that compared to younger adults, older adults had significantly higher entropy in the sensorimotor cortex, basal ganglia, hippocampus, thalamus, and occipital lobe. Higher entropy in many of these regions was associated with worse physical function and higher body mass index in older adults. Findings suggest that spatial entropy in brain network landscapes may be a marker of declining physical function. Modifiable factors, such as losing excess weight, may help to ameliorate aging-related brain entropy increases in regions such as the sensorimotor cortex, which may in turn help to preserve physical function in older adults.

## Introduction

The number of Americans aged 65 or older is expected to increase by 23 million people from 2025 to 2055 (*The Demographic Outlook: 2025 to 2055*, 2025). The demographic shift is will be accompanied by an increased burden of aging-related disease on healthcare systems (Vincent & Velkoff, 2010). Advanced age is the strongest predictor of physical function decline (Ferrucci et al., 2016; Ko et al., 2010; Layne et al., 2017), which is associated with lower quality of life (Petnehazy et al., 2024), higher risk of falls or injury (O’Hoski et al., 2020; Smee et al., 2012), and higher mortality (Cesari et al., 2009; De Buyser et al., 2013; Majer et al., 2011).

In addition to chronological age, physical function is influenced by factors such as overweight/obesity, sedentary lifestyle, and diet quality (Robinson et al., 2013), which contribute to aging pathology through many cellular and molecular mechanisms (Gladyshev et al., 2021). Together, these mechanisms reflect the principle that biological systems accumulate damage over time. Damage can be conceptualized as “disorder” (Meyer et al., 2025) quantified by Shannon entropy (Shannon, 1948).

The observation that biological systems tend to become more disordered with age has led several to point out parallels with the 2^nd^ Law of Thermodynamics (Cummings et al., 2025; Drachman, 2006; Strehler, 1962), which dictates that the entropy of a system must increase over time. Conceptualizing aging as a potentially ameliorable increase in entropy has gained attention in recent years, but primarily in the context of microscopic biological systems. To what extent the increases in entropy extend to large scale functional organization in the human body is an open question.

The brain is one large scale system that is of particular interest in aging and to which the concept of increasing entropy may be particularly appropriate (Drachman, 2006). The brain is often modeled as a network of regions tied together by functional connections (Biswal et al., 1995; Bullmore & Sporns, 2009). Network neuroscience has led to identification of distinct communities within brain networks that associate with specific cognitive and physical functions (Power et al., 2011; Yeo et al., 2011). When mapped into the brain together, these communities form functional “community landscapes”, which serve as maps of cognitive and physical function in the brain. Physical function has been shown to be related to the spatial organization of the sensorimotor network in older adults (Hugenschmidt et al., 2014; Laurienti et al., 2023). However, prior investigations required *a priori* specification of expected, “ideal” community structure and did not specifically assess the spatial disorder of the networks.

Recently, methodology has been developed to analyze the spatial disorder in community landscapes across the whole brain in a completely data-driven manner (McIntyre et al., 2025). This methodology utilizes a version of Shannon entropy that quantifies the spatial entropy of brain networks such that regions with spatially ordered communities have low entropy and regions with spatially disordered communities have high entropy. Entropy is quantified for each voxel of brain network images resulting in spatial entropy maps for each participant.

In this work, we apply novel methodology for mapping brain network spatial entropy to a cohort of cognitively normal, community-dwelling older adults (n = 192) from the Brain Networks and Mobility (B-NET) study. We also examined brain entropy in a comparison cohort of healthy younger adults (n = 30). We compared the brain entropy maps of older and younger adults, hypothesizing that older adults would have higher entropy than younger adults across the whole brain. We also examined the relationship between physical function, measured by the Expanded Short Physical Performance Battery (eSPPB), and brain network entropy in older adults. Based on prior work (Hugenschmidt et al., 2014; Laurienti et al., 2023), we hypothesized that poorer physical function would be associated with higher entropy (more spatial disorder) in the sensorimotor network. Finally, in exploratory analyses, we assessed the relationships between brain entropy and several markers of general health to generate hypotheses about how brain entropy might be modified in future studies to improve aging outcomes.

## Results

### Participant Characteristics

Participants in the cohort of older adults (n = 192, 108 female) had an average age of 76.4 years (standard deviation = 4.7 years) with average eSPPB scores of 2.0 (standard deviation = 0.5), and average body mass index (BMI) of 28.4 kg/m^2^ (standard deviation = 5.5 kg/m^2^).

In the younger cohort (n = 30, 19 female), participants had an average age of 30.0 years (standard deviation = 3.8 years) with average eSPPB scores of 2.8 (standard deviation = 0.2) and average BMI of 25.5 kg/m^2^ (standard deviation = 5.1 kg/m^2^).

Welch’s two-sample T-tests were used to assess whether these measures differed between the younger and older cohort. Relative to the young cohort, the older cohort had worse average eSPPB scores (*p* < 0.001) and significantly higher average BMI (*p* = 0.007). More details on general physical and cognitive health measures in these two samples are presented in Supplementary Table 1 and in previous work from the B-NET study (Laurienti et al., 2023; Thompson et al., 2023). For information about the correlation of key demographic and biological variables in the older adults, see Supplementary Figure 1.

### Brain Network Entropy – Comparing Younger and Older Adults

Spatial entropy maps were generated for the brains of each older and younger adult participant. In Figure 1, we show a representative participant’s community landscape and corresponding entropy maps along with the average entropy map for the older and younger cohorts.

**Figure 1.**
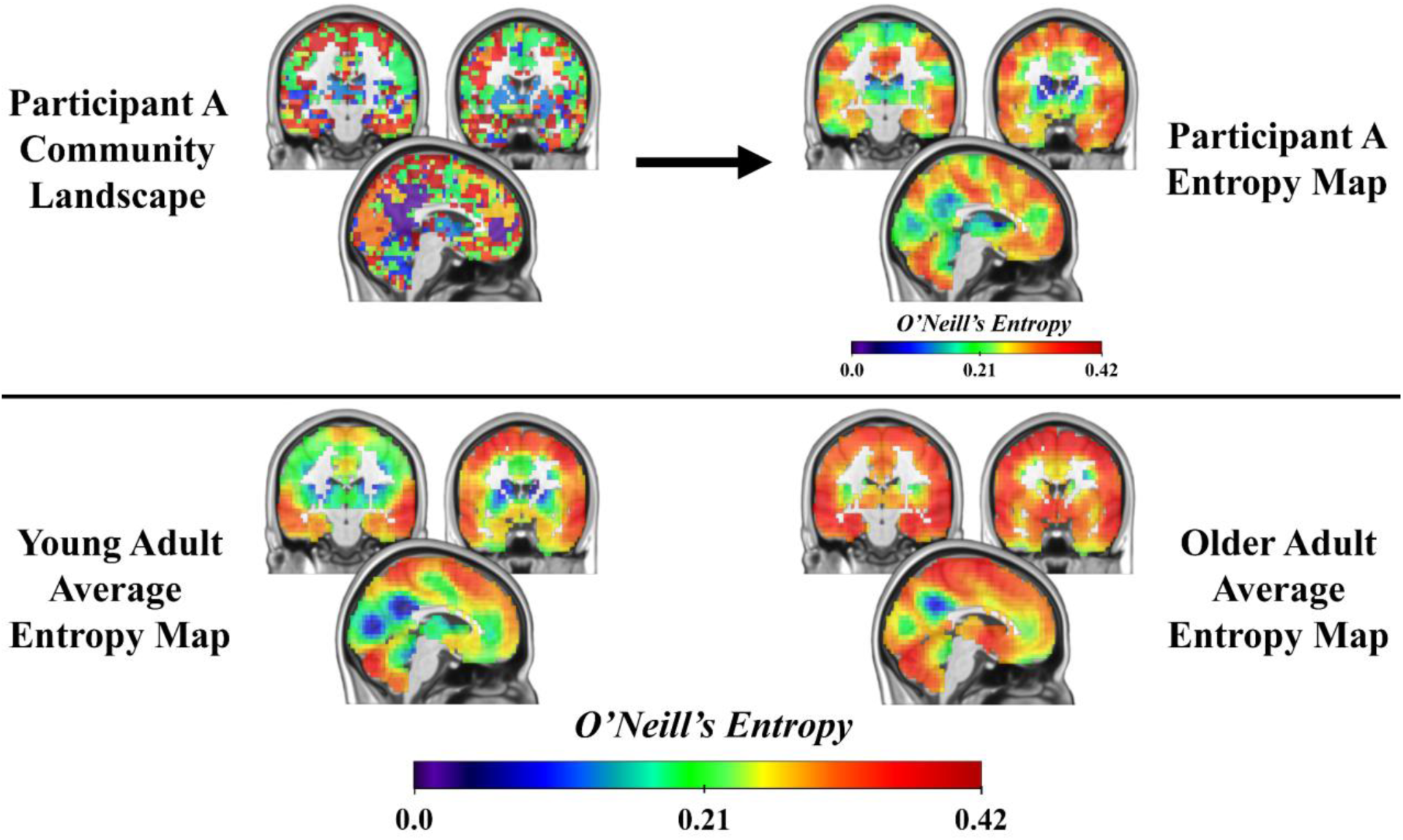
A representative community landscape and corresponding entropy map from young adult participant “A” is shown along with group-average entropy maps for the younger and older cohorts. Regions with higher entropy tend to have very spatially disordered community representation in nearby voxels whereas regions with lower entropy tend to have more spatially ordered communities. The MNI coordinates of the slices used in this figure are X = - 5, Y = -19, Y = 0.

To assess statistical differences in entropy maps between the older adult cohort and the younger adult cohort, spatial entropy maps were compared using two-sample t-tests. Figure 2 shows maps of T-values from the comparison. Images from both thresholded and unthresholded maps are shown to both highlight statistically significant differences while also allowing patterns from the whole brain to be observed (Misic, 2025). Warm colors indicate regions with higher entropy in older adults relative to younger adults. In older adults, spatial entropy was significantly higher in regions directly related to physical function, including the sensorimotor cortex, basal ganglia, and thalamus, as well as in the hippocampus and occipital lobe. There were no regions with significantly higher spatial entropy in younger adults compared to older adults.

**Figure 2.**
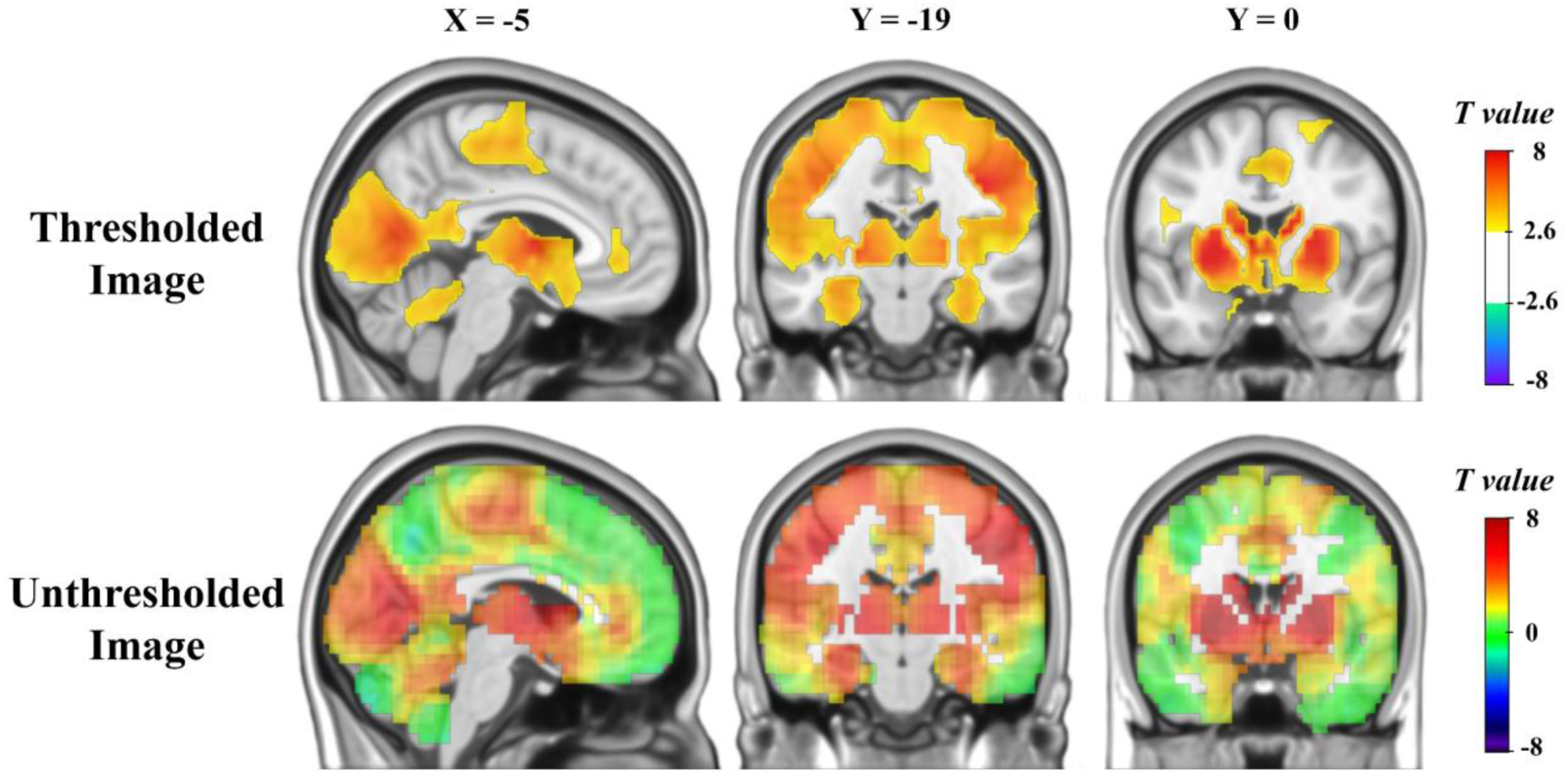
Thresholded and unthresholded maps of T-values from a two-sample t-test comparing entropy maps of a cohort of older adults and a cohort of younger adults. Warmer colors indicate regions with higher entropy in the older adults.

### Brain Network Entropy - Physical Function in Older Adults

To determine whether physical function was associated with brain network spatial entropy, eSPPB scores from the older participants were regressed against entropy maps. Figure 3 shows maps of T-values from the regression. There was a negative association between eSPPB and spatial entropy. Specifically, poorer performance on the eSPPB was associated with higher entropy in regions including the sensorimotor cortex, basal ganglia, and bilateral middle frontal gyri. Better eSPPB was not significantly associated with higher entropy in any brain regions.

**Figure 3.**
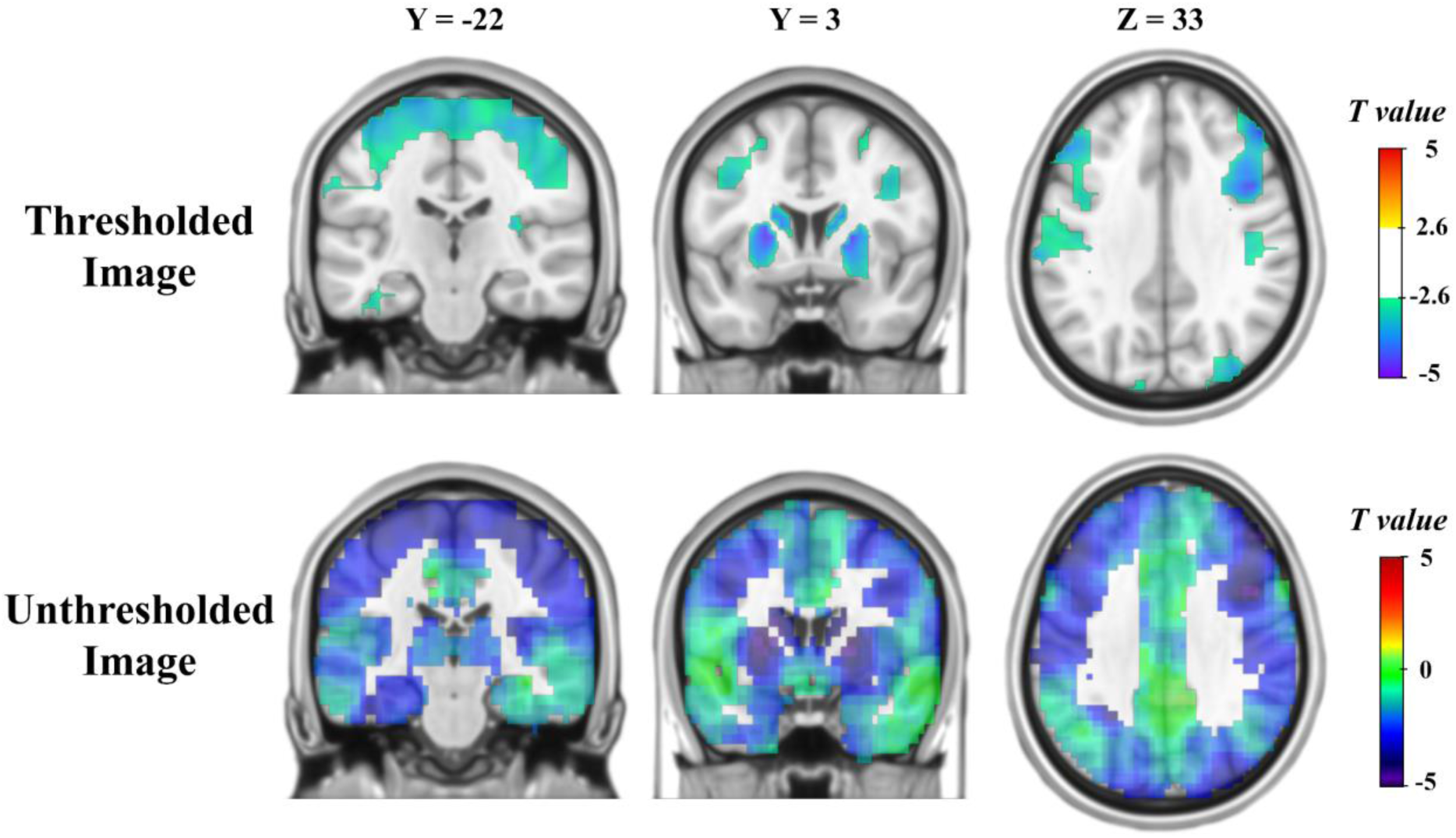
Thresholded and unthresholded maps of T-values from a regression of eSPPB against brain entropy maps in the cohort of older adults. Cooler colors indicate a negative association between eSPPB and entropy.

### Brain Network Entropy – Additional Health Factors in Older Adults

Previous work from the B-NET study found that BMI was related to community structure consistency in the sensorimotor network (Laurienti et al., 2023). Therefore, we evaluated whether BMI was related to brain entropy maps. A regression model was fit with BMI as the independent variable of interest and entropy maps as the outcome. Figure 4 shows maps of T-values from the regression, which indicated that BMI was positively associated with spatial entropy. Higher BMI was associated with higher entropy in portions of every cortical lobe with the strongest associations being in the left sensorimotor cortex and the bilateral posterior insula. Higher BMI was not significantly associated with lower entropy in any brain regions.

**Figure 4.**
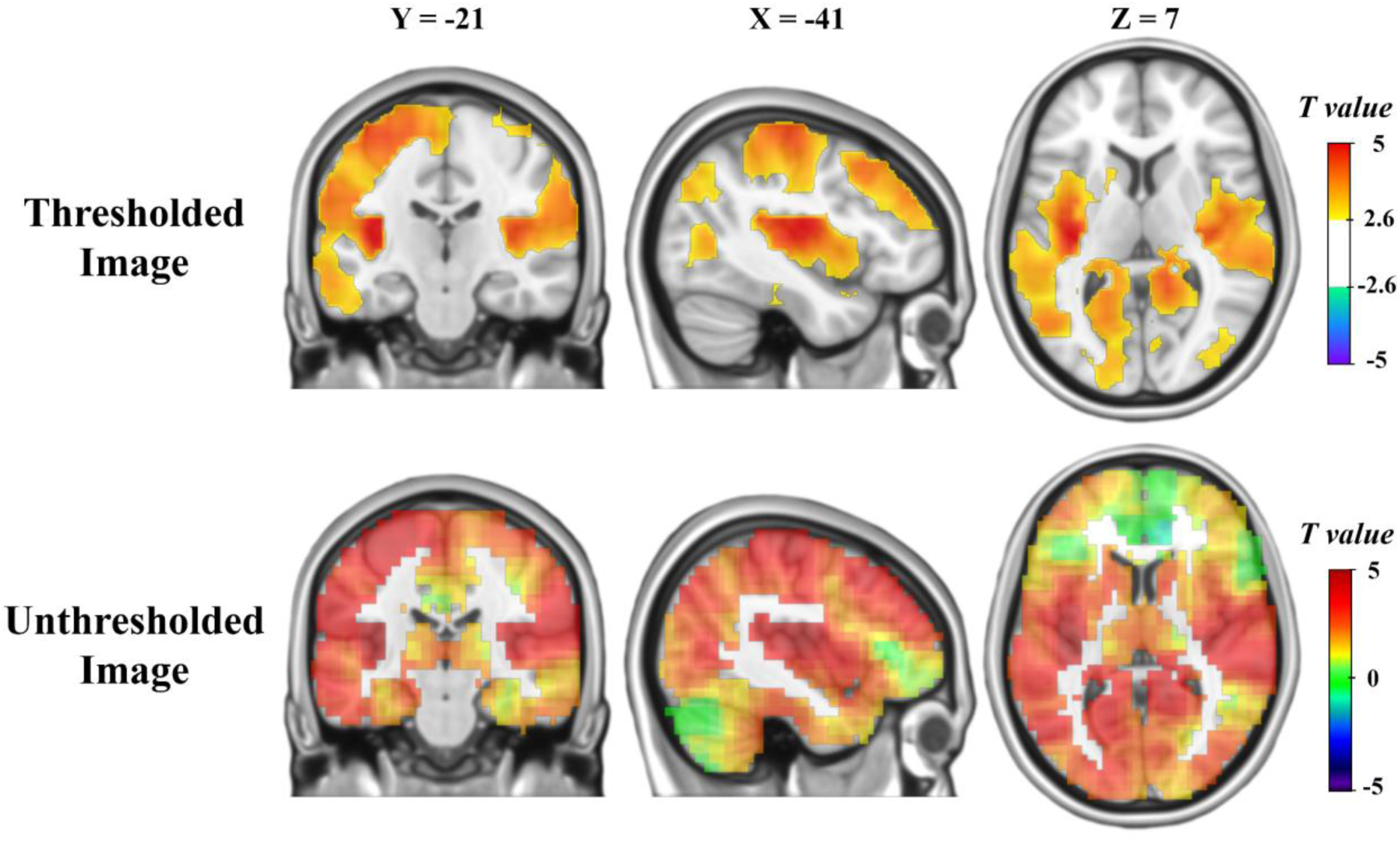
Thresholded and unthresholded maps of T-values from a regression of BMI against brain entropy maps in the cohort of older adults. Warmer colors indicate a positive association between BMI and entropy.

A young adult may eventually be an older adult with a healthy weight, or they may eventually be an older adult with obesity. Based on Figure 2, we expect brain entropy to be higher in older adults. Based on Figure 4, we expect brain entropy to be highest in older adults with obesity. We have not yet assessed whether older adults with a healthy weight have higher brain entropy than younger adults, or whether elevated brain entropy is unique to older adults with obesity. In Figure 5, we show the result of two two-sample t-tests comparing our cohort of younger adults to older adults (n = 52) with “normal” weight (BMI between 18.5 and 25) as well as a comparison of younger adults to older adults (n = 61) with obesity (BMI ≥ 30). Older adults with normal weight had higher entropy than younger adults, particularly in the basal ganglia, sensorimotor cortex, thalamus, and occipital lobe. The difference in entropy between younger adults and older adults with obesity followed a similar spatial pattern but was substantially more pronounced and widespread in the brain.

**Figure 5.**
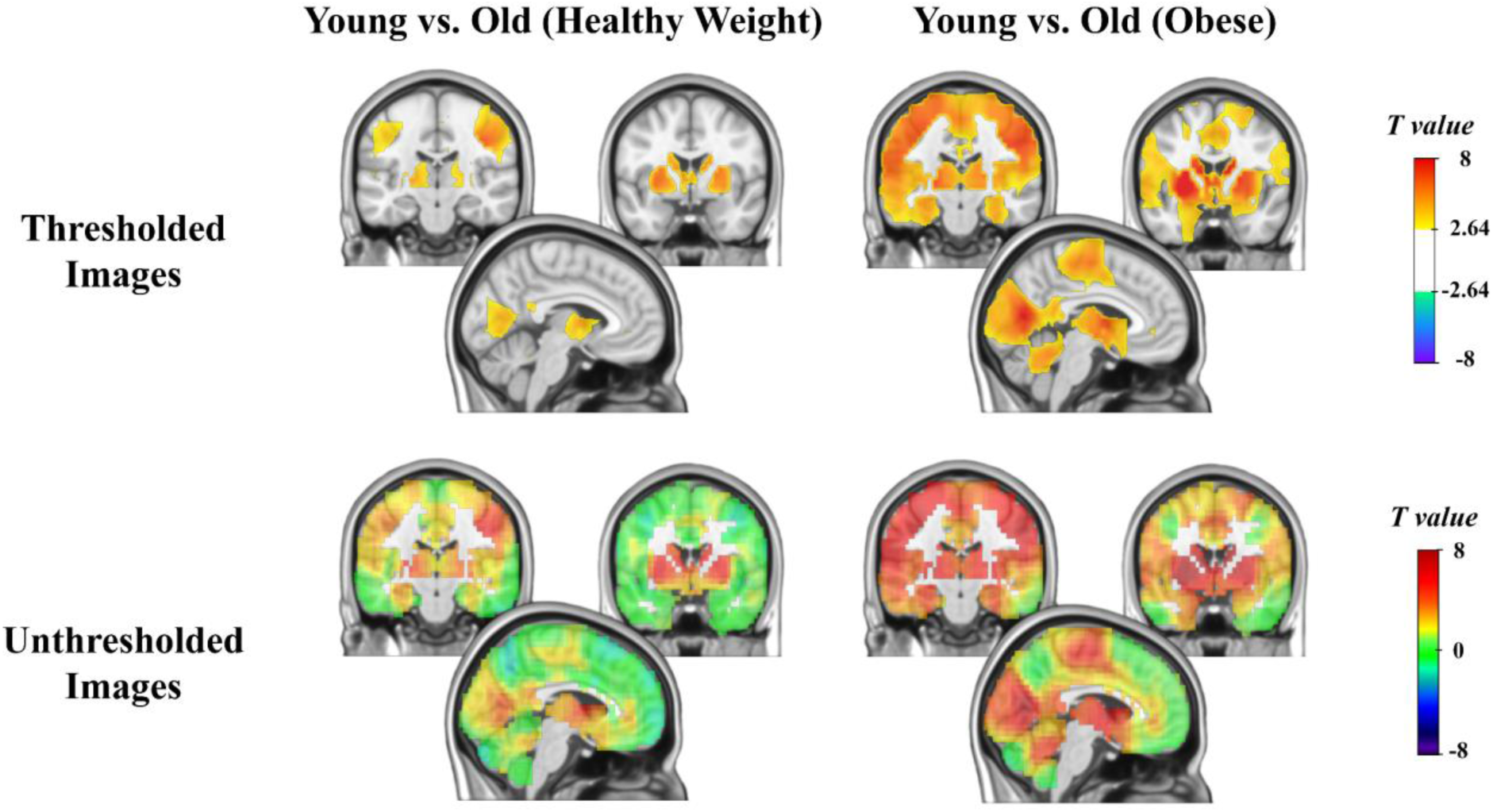
Thresholded and unthresholded maps of T-values from two-sample T-tests comparing a) younger adults to older adults with normal weight, and b) younger adults to older adults with obesity. Warmer colors indicate higher entropy in the older group. The MNI coordinates of the slices used in this figure are X = -5, Y = -19, Y = 0.

## Discussion

We showed that brain entropy was higher in older adults than in younger adults, particularly in the sensorimotor cortex, basal ganglia, thalamus, hippocampus, and occipital lobe. Higher entropy in regions including the sensorimotor cortex, basal ganglia, and middle frontal gyrus was associated with worse physical function in older adults. Finally, BMI was associated with higher entropy in regions including the sensorimotor cortex, occipital lobe, and posterior insula. We interpret findings as evidence that spatial entropy in resting-state brain networks may be a marker of poor physical function. Modifiable factors, such as losing excess weight, may help to ameliorate age-related brain entropy increases in regions associated with sensory and motor processing, which may in turn protect physical function.

The finding that entropy was higher in the brains of older adults compared to younger adults supported our hypothesis. Regions that had the most pronounced difference in entropy included regions associated with motor control, such as the motor cortex, basal ganglia, and cerebellum, as well as regions associated with lower order sensory processing, such as the visual cortex. Based on prior work (McIntyre et al., 2025), we expect that higher entropy in a brain region should be associated with deficits in the physical and cognitive functions for which that region is most directly responsible. This view is consistent with the finding that older adults have higher entropy in lower order motor and visual processing regions as both physical function and visual ability decline with aging.

In addition to motor and sensory regions, we also observed higher entropy in the hippocampus of older adults compared to younger adults. The hippocampus has long been known for its role in memory formation, learning, and spatial awareness (Bird & Burgess, 2008; Olton et al., 1979; Scoville & Milner, 1957). Higher entropy in this region in older adults may reflect a wide range of cognitive deficits. The older adults did have significantly worse cognitive function than the younger adults across several cognitive domains (Supplementary Table 1). The relationship with the hippocampus is also particularly fascinating given extensive work suggesting that this structure is one of the first to be affected by aging-related diseases such as Alzheimer’s disease (Barnes et al., 2009; Rao et al., 2022).

As hypothesized, we also observed that poorer physical function was associated with higher entropy in cortical regions that are part of the sensorimotor network. Poorer physical function was also associated with higher entropy in the basal ganglia. The basal ganglia finding was not specifically hypothesized but was unsurprising given that these areas are known to be crucial for modulatory motor control and motor planning (Arber & Costa, 2022; Dhawale et al., 2021; Groenewegen, 2003; Young et al., 2025). The relationship between physical function and basal ganglia community organization was not observed in prior work (Hugenschmidt et al., 2014; Laurienti et al., 2023) that were not able to do whole-brain analyses (like those performed here) due to methodological limitations. The basal ganglia finding is very intuitive and highlights the capacity for entropy-mapping methodology to identify disordered community structure in the whole brain without biasing analyses towards recreating prior work.

Additionally, we observed that poorer physical function was associated with higher entropy in the bilateral middle frontal gyri. These regions are typically associated with cognitive functions (Briggs et al., 2021; Marek & Dosenbach, 2018; Menon, 2011; Menon & D’Esposito, 2022) rather than motor function, so we did not hypothesize this relationship. We note that several previous studies have identified relationships between physical and cognitive decline in older adults (Auyeung et al., 2008; Binder et al., 1999; Moritz et al., 1995; Njegovan et al., 2001). Speculatively, impaired function in this region may result in difficulty adapting to environmental changes, which may result in hesitant, uncoordinated movements. The middle frontal gyrus is routinely targeted using techniques such as transcranial direct current stimulation, which may improve physical function in older adults (Lo et al., 2025). Future work may assess whether stimulating the middle frontal gyrus reduces local brain entropy, and whether any entropy reduction is accompanied by lasting improvements in physical function.

The finding that BMI is strongly related to higher brain entropy – including in regions associated with physical function – suggests that it may be worthwhile to test whether behavioral and/or pharmacological interventions modify brain entropy. For example, glucagon-like peptide-1 (GLP-1) analogs are increasingly popular for weight loss, though there is still debate regarding the benefits and risks of prescribing these drugs in older adults (Azzolino & Lucchi, 2025; Dadwani et al., 2023; Prokopidis et al., 2025). Future work may benefit from considering how these drugs and/or behavioral interventions impact brain entropy in older adults.

This study acknowledges several limitations. Because all data shown in this work is cross-sectional, causality in relationships between variables of interest and brain entropy cannot be determined. Future work investigating targeted interventions will help establish directionality in these relationships. Additionally, this work was largely focused on the relationship between brain entropy and physical function. Future work should explore the relationship between brain entropy and other measures of healthy aging, such as cognitive function.

The motivation for studying the brain as a complex network stems from the intuition that it should be considered as a gestalt - physical function is not entirely controlled by any one brain region, nor is any one region exclusively involved in physical function (Pessoa, 2014). Spatial entropy mapping gives us a way to quantify the brain’s local organization within the context of the system as a whole. In this work, we use spatial entropy to show – for the first time – that aging is associated with brain network community structure deterioration, and that disordered community structure in the sensorimotor cortex, basal ganglia, and other regions is associated with poorer physical function. This finding gives us tangible markers of physical function in the brain. Future work will now be able to tailor interventions towards modifying entropy in these regions to improve aging outcomes.

## Methods

### Study Design and Sample

All data in this work was collected by the Brain Networks and Mobility (B-NET) study. B-NET was a longitudinal, observational study of cognitively normal community-dwelling older adults (n=192) aged 70 and older. The present work includes data only from baseline study visits. At baseline, participants provided extensive information about their health history, completed physical function and cognitive testing, and completed brain MRI scanning. In addition to the large cohort of older adults, a smaller cohort (n=30) of younger adults aged 22-36 years completed baseline study activities so that measures from the older adults could be compared to a young, healthy sample.

Participants were recruited from direct mailings, flyers, word of mouth, and the community newsletter from the Sticht Center for Healthy Aging and Alzheimer’s Prevention at Wake Forest University School of Medicine. Exclusion criteria included being a single or double amputee, having musculoskeletal implants severe enough to impede physical function testing, being dependent on a walker or another person to ambulate, having undergone surgery in the past 6 months, having any serious or uncontrolled chronic disease, having major uncorrected hearing or vision problems, plans to relocate in the two years following baseline study visit, or active participation in a behavioral intervention trial. Neurological/psychiatric exclusion criteria included diagnosis of a neurologic disease that affected mobility, prior traumatic brain injury with residual deficits, history of brain tumor, seizures within the last year, diagnosis of bipolar disorder, schizophrenia, any other psychotic disorder, hazardous alcohol use (>21 drinks per week), or evidence of impaired cognitive function. Cognitive impairment was assessed according to scores on the Montreal Cognitive Assessment (MoCA). Participants with Montreal Cognitive Assessment scores of 20 or lower were excluded. Scores of 21-25 were reviewed by the B-NET study neuropsychologist to determine eligibility. Finally, participants were excluded if they were unable or unwilling to complete brain MRI scanning.

All participants gave written informed consent to participate in the activities associated with this study as approved by the Institutional Review Board (IRB) of the Wake Forest University School of Medicine (IRB protocol #IRB00046460; approval date: August 27, 2020).

### Study Measures

Baseline measures from the B-NET study were collected over the course of two study visits that were typically completed within one month of each other. Most measures were collected in the first visit. The second visit was for MRI scanning. The only demographic variable used for analyses in this work was age. Height and weight were measured using a wall-mounted stadiometer and calibrated scale. These measures were then used to compute each participant’s body mass index (BMI). Additional measures were included in supplemental exploratory analyses – for more details, see the Extended Methods and Results in the Supplementary Material.

Physical function was quantified by the expanded short physical performance battery (eSPPB)(Simonsick et al., 2001). The eSPPB was adapted from the original SPPB (Guralnik et al., 1994) in order to address ceiling effects that limit the utility of SPPB score in cohorts of well-functioning older adults, such as the B-NET sample. The eSPPB is composed of four components that measure different dimensions of physical function. The first component, balance, is measured by asking participants to hold a side-by-side posture for 10 seconds, followed by the semi-tandem, tandem, and one-leg positions for 30 seconds each. The second component, gait speed, assesses the participant’s 4-meter gait speed (m/s). The third component, complex gait speed, repeats the second component test but requires participants to complete narrow steps within two parallel lines spaced 20 cm apart. The fourth component, lower extremity strength, assesses the time it takes participants to stand up from a seated position five times. Each component is individually scored from 0 to 1 according to the best possible performance for each component. The sum of the component scores is the eSPPB total score, which is continuous ranging from 0 to 4. Two participants had missing data that prevented calculating their eSPPB score. In this work, analyses that included eSPPB as a variable were restricted to the 190 older adults with eSPPB scores. All 30 younger adults had sufficient data to quantify their eSPPB scores.

### Brain Image Acquisition

Prior work from the B-NET study provides a detailed description of all brain imaging completed at B-NET imaging visits (Laurienti et al., 2023). Here, we summarize only the imaging acquisition relevant to analyses for the present work. All brain images were collected on a Siemens 3T Skyra MRI scanner equipped with a 32-channel head coil. Each scan session lasted approximately one hour. An anatomical image was collected using T1-weighted 3D volumetric MPRAGE sequence (TR = 2000ms, TE = 2.98 ms, number of slices = 192, 1.0mm isotropic voxels, FOV = 256mm, scan duration = 312s). Resting-state functional MRI data were collected using a blood oxygenation level-dependent (BOLD) (Ogawa et al., 1990) weighted echo planar imaging (EPI) sequence (TR = 2000ms, TE = 25ms, number of slices = 35, voxel dimension = 4.0×4.0×5.0mm, FOV = 256mm, duration = 7m14s). During resting-state scans, participants viewed a black fixation cross displayed on a white background.

### Image Preprocessing

Statistical Parametric Mapping version 12 (SPM12, https://www.fil.ion.ucl.ac.uk/spm/software/spm12/) was used for structural image segmentation. Gray matter and white matter segmented images were summed, and any voxel with at least 0.5 probability was retained as brain parenchyma. Non-brain tissue and cerebrospinal fluid (CSF) were excluded from brain masks. Structural images were masked, visually inspected and manually cleaned to remove any remaining non-parenchymal tissues using MRIcron software (https://www.nitrc.org/projects/mricron). Two observers manually checked masked images to ensure accurate full-brain coverage. The masked, cleaned T1-weighted images were spatially normalized to the Montreal Neurological Institute (MNI) template using Advanced Normalization Tools (ANTs, https://antsx.github.io/ANTs/).

BOLD-weighted images underwent distortion correction using FMRIB’s “Topup” Software Library (FSL, https://fsl.fmrib.ox.ac.uk/fsl/docs/<u>#/</u>). The first ten volumes of the BOLD images were removed to allow for signal normalization. Slice time correction and realignment of the functional images was completed using SPM12. BOLD images were coregistered to native-space anatomical images, then warped to MNI space using the transformation derived from ANTs. Data were band-pass filtered (0.009 – 0.08 Hz) to account for low-frequency drift and physiological noise. Confounding signals were regressed out from the filtered data; this included global average signal from white matter, gray matter, CSF, and the 6 rigid-body motion parameters that are obtained during the realignment process. Finally, motion correction was applied using the motion scrubbing procedure (Power et al., 2014), which involves removing any volumes that contain both excessive (>0.5 mm) movement and excessive signal change (>0.5 DV_GM_).

### Brain Network Generation

Functional brain networks were generated by first finding the Pearson correlation of the BOLD signal time series for every pairing of voxels in each participant’s brain network. The result was an *N*x*N* correlation matrix where *N* is the number of network nodes (*N* ≅ 20,000). This correlation matrix was converted to an adjacency matrix *A_ij_* by thresholding connections to retain only the strongest edges that satisfied the equation *S =* log(*N*) / log(*k*), where *k* is the average number of retained connections per node and *S* was set to 2.5 according to previous work (Hayasaka & Laurienti, 2010; Laurienti et al., 2023). Retained connections were set to 1, resulting in a sparse, binarized adjacency matrix for each participant. This adjacency matrix served as the data structure for each participant’s brain network. The end result of this process is a brain network for every participant where nodes represent voxels and edges represent strongly synchronized BOLD signal between two voxels.

### Network Community Detection

A network community is a group of nodes that are more connected to each other than they are to nodes from other network communities (Newman, 2006; Newman & Girvan, 2004). Communities were identified in each participant’s brain network such that each voxel (node) was assigned to a single community. Community partitioning was optimized using the modularity metric *Q* (Newman, 2006; Newman & Girvan, 2004). A dynamic Markov process (Delvenne et al., 2010) was used to identify the network community partitioning that maximized *Q*. As this algorithm contains a stochastic component, it was run 100 times, and the partitioning that yielded the highest *Q* value for a given network was used to represent the categorical community structure of that network. The community assignment of each voxel (node) can be mapped into brain space to serve as a functional brain network landscape, illustrating brain regions that have highly diverse or homogenous functional community affiliations.

### Spatial Entropy Calculation

Spatial entropy was quantified for each voxel in each participant’s brain network according to novel methodology that allows quantification of the local spatial disorder of brain network landscapes (McIntyre et al., 2025). For each voxel *i*, a spatial “neighborhood” encompassing all voxels within three voxels from *i* was identified. The neighborhood spatial entropy was then quantified using a modified version of O’Neill’s entropy *H_O_* (O’Neill et al., 1988), which is an adaptation of Shannon’s entropy (Shannon, 1948) developed specifically for spatial analyses. *H_O_* was calculated for each voxel neighborhood as

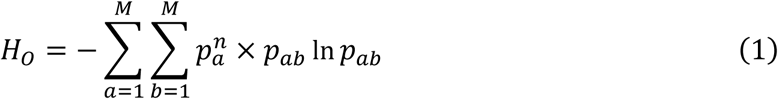

where *M* is the number of communities in the network, *p_a_^n^* is the proportion of voxels in the whole brain network belonging to community *a*, and *p_ab_* is the proportion of contiguous (touching) voxels with one voxel belonging to community *a* and the other belonging to community *b*. The *p_ab_*ln(*p_ab_*) term quantifies the heterogeneity of adjacent pairings of voxels within a neighborhood. *H_O_* is highest when each combination of contiguous pairings of communities are equally represented. *H_O_* is lowest when only one module is present in the neighborhood (and, therefore, all two-voxel pairings represent the same combination of communities). The *p_a_^n^* term adjusts the contributions of voxels from community *a* to the overall entropy of the neighborhood such that voxels from larger communities are weighted more than voxels from smaller communities. The reason for including this term is that without it, the presence of voxels from very small communities in a neighborhood can inappropriately raise the minimum theoretical entropy of the neighborhood (McIntyre et al., 2025). The *H_O_* value of the neighborhood was assigned to voxel *i*. The result of carrying out this procedure for each voxel in each participant’s brain network was a map of each participant’s brain network spatial entropy.

### Statistical Analysis of Entropy Maps

SPM12 was used for statistical analyses of entropy maps. SPM two-sample t-tests were used to compare the entropy maps of the cohort of older adults with the cohort of young adults. In supplemental analyses, an additional model was fit adjusting for the number of volumes removed from functional scans due to in-scanner motion.

The primary motivation of the B-NET study was to assess the relationship between brain networks and physical function in older adults. Therefore, within the older sample, we used eSPPB scores as a regressor to identify whether physical function was related to brain entropy maps. In supplemental analyses, we fit the same regression model but with the number of volumes removed from scan due to in-scanner motion being included as a covariate. Because prior work in this sample has found that BMI is associated with the spatial consistency of the sensorimotor subnetwork (Laurienti et al., 2023), we also assessed whether BMI was associated with entropy maps. Again, we fit both an unadjusted regression model and a model adjusting for in-scanner motion.

In additional exploratory analyses, we used regression models to determine whether age was associated with entropy maps in older adults, and whether other measures of general health - type 2 diabetes history, hypertension history, and physical activity – were related to brain entropy.

For all imaging analyses, significance at the voxel level was set at *p* < 0.005. To correct for multiple comparisons, we further required voxel cluster extent to be significant at *p* < 0.05. We show both thresholded (i.e., voxels that are statistically significant following cluster correction) and unthresholded T-value maps that result from SPM t-tests and regressions to maximize transparency and encourage contemplation of the spatial patterns of observed relationships without being bound to significance thresholds (Misic, 2025).

## Supporting information

Extended Methods and Results

## Notes

**Conflict of interest:** The authors declare no conflicts of interest.

### Competing Interest Statement

The authors have declared no competing interest.

