## Extended Methods and Results for "Poorer physical function is associated with elevated spatial entropy in the aging brain network landscape"

*Additional Health and Behavioral Measures*

All additional health measures are based on assessments performed at the participants' baseline visits. Diabetes was defined as anyone that is currently taking diabetes medication or that had fasting serum glucose > 130 mg/dL. Hypertension was defined as anyone with systolic blood pressure > 150 mmHg or anyone taking hypertension medication with systolic blood pressure > 130 mmHg. Physical activity was assessed using responses to the Community Healthy Activities Model Program for Seniors (CHAMPS) questionnaire (Stewart et al., 2001). Responses were binarized such that physical activity was coded as "1" if the participant reported engaging in at least 150 minutes of moderate physical activity per week and "0" if they did not.

### Extended Results

*Supplementary Table 1. Variable Differences between Younger and Older Adults*

| Variables | Older Adults (n = 192*) | Younger Adults (n = 30) | *p*-value |
|---|---|---|---|
| Age (years) | 76.4±4.7 | 30.0±3.8 | **<0.0001** |
| Expanded Short Physical Performance Battery | 2.0±0.5 | 2.8±0.2 | **<0.0001** |
| Body Mass Index (kg/m$^2$) | 28.4±5.5 | 25.5±5.1 | **0.0071** |
| Systolic Blood Pressure (mmHg) | 138.7±17.9 | 118.2±17.3 | **<0.0001** |
| Fasting Glucose (mg/dL) | 106.3±23.0 | 89.4±7.1 | **<0.0001** |
| Montreal Cognitive Assessment Raw Total Score | 25.5±2.2 | 28.0±2.0 | **<0.0001** |
| Digit Symbol Substitution Task Score | 55.2±12.2 | 84.9±12.8 | **<0.0001** |
| Trailmaking A Time (seconds) | 36.8±11.1 | 23.8±8.4 | **<0.0001** |
| Trailmaking B Time (seconds) | 98.7±44.0 | 53.1±15.3 | **<0.0001** |
| fMRI – Volumes Removed Due to Motion | 7.9±13.9 | 8.4±9.8 | 0.813 |

Cohort mean values for key variables from analyses. Values following '±' indicate the standard deviation of the variable in the cohort. Statistical comparison between cohorts was carried out using Welch's T-tests. *For some measures, some participants did not have data available (Expanded Short Physical Performance Battery had 190 of 192 older adults with data available, Trailmaking B had 191 of 192 older adults with data available). In these cases, statistical comparison with the younger cohort was based on those older adults who had data for the measure.

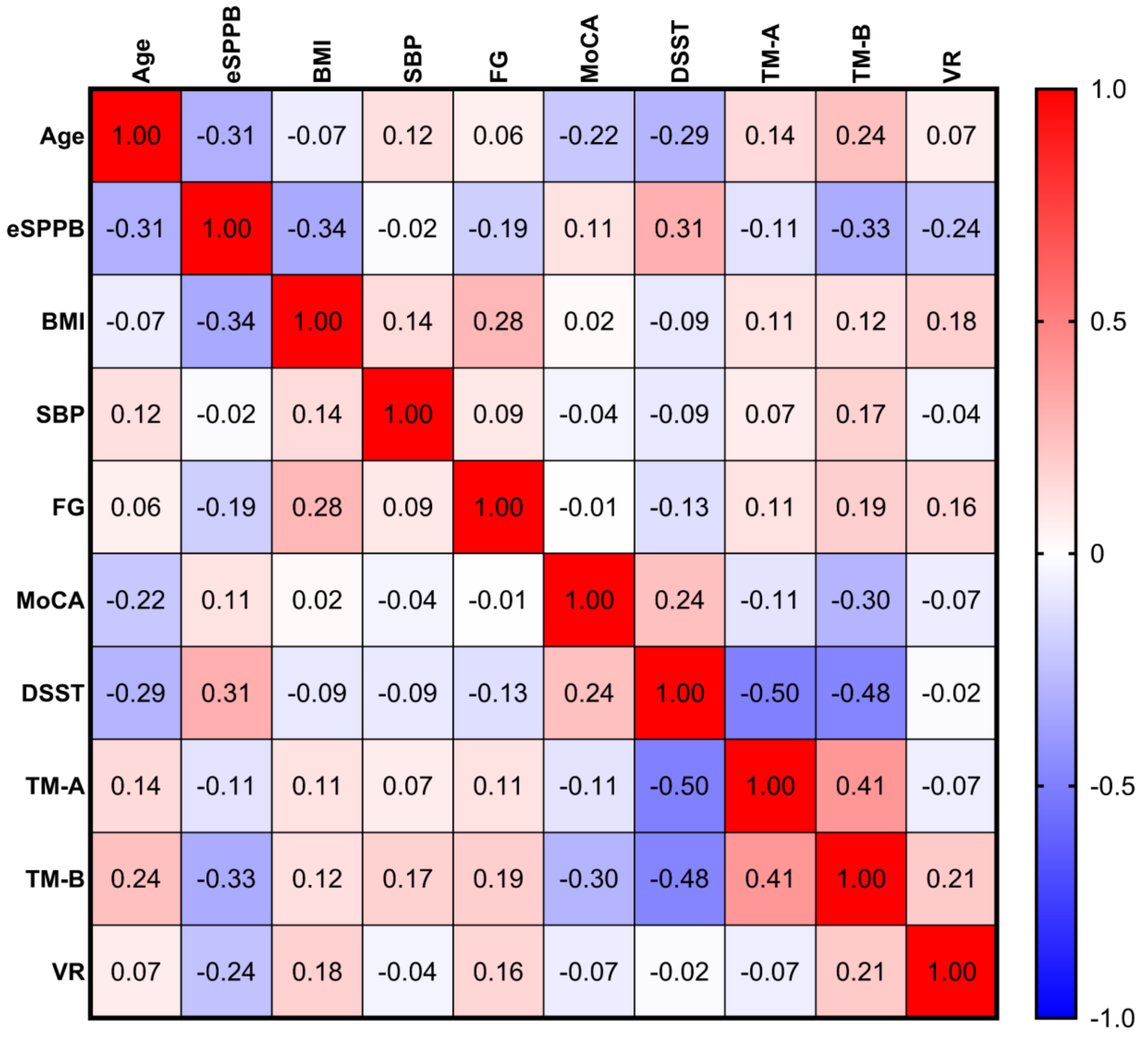


**Supplementary Figure 1.** Spearman correlation matrix for key variables in the cohort of older adults. Red indicates positive correlation while blue indicates negative correlation. As with Supplementary Table 1, correlations are based only on participants with data available. The variables follow the same order that is presented in Supplementary Table 1. Numeric values represent Spearman's ρ from the relationship between variables.

*eSPPB* – Expanded Short Physical Performance Battery

*BMI* – Body Mass Index

*SBP* – Systolic Blood Pressure

*FG* – Fasting Blood Glucose

*MoCA* – Montreal Cognitive Assessment Raw Total Score

*DSST* – Digit Symbol Substitution Task Score

*TM-A* – Trailmaking A Time

*TM-B* – Trailmaking B Time

*VR* – Volumes Removed from resting-state fMRI scans due to motion

Supplementary Figure 1 shows the Spearman correlation coefficient between key variables in analyses and some additional variables of interest, such as performance on cognitive tests, for the sample of older adults. We note that some key variables of interest, such as eSPPB and BMI, were interrelated. These variables also shared a common association with in-scanner motion.

Supplementary Figure 2 shows results from a t-test of the entropy maps of younger and older adults with adjustment for the number of volumes removed from scans due to motion. Results did not meaningfully change from Figure 1 of the main text.

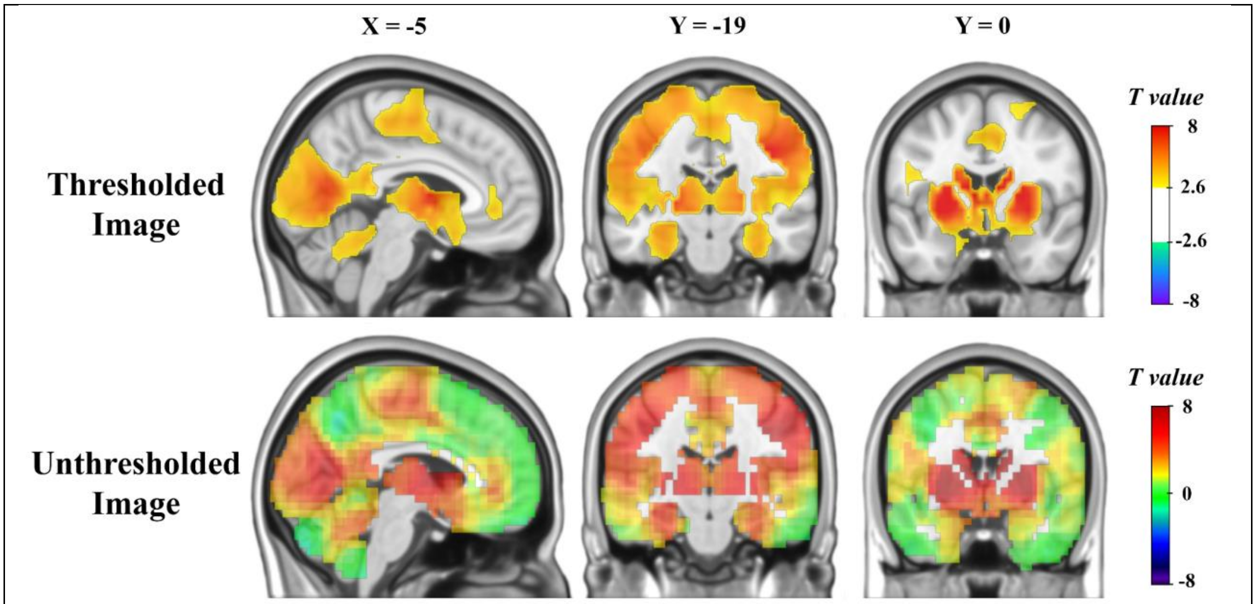


**Supplementary Figure 2.** Thresholded and unthresholded maps of T-values from a two-sample t-test comparing entropy maps of a cohort of older adults and a cohort of younger adults. Warmer colors indicate regions with higher entropy in the older adults. Cooler colors indicate regions with higher entropy in the younger adults. The t-test controlled for the number of volumes removed from scans to correct for head motion.

Supplementary Figure 3 shows results from a regression of eSPPB against entropy maps while adjusting for in-scanner motion. Results were similar to Figure 2 from the main paper. The most noticeable difference was that the association in the basal ganglia was no longer significant following cluster correction for multiple comparisons, though the unthresholded image indicates that the effect estimate was only slightly weakened.

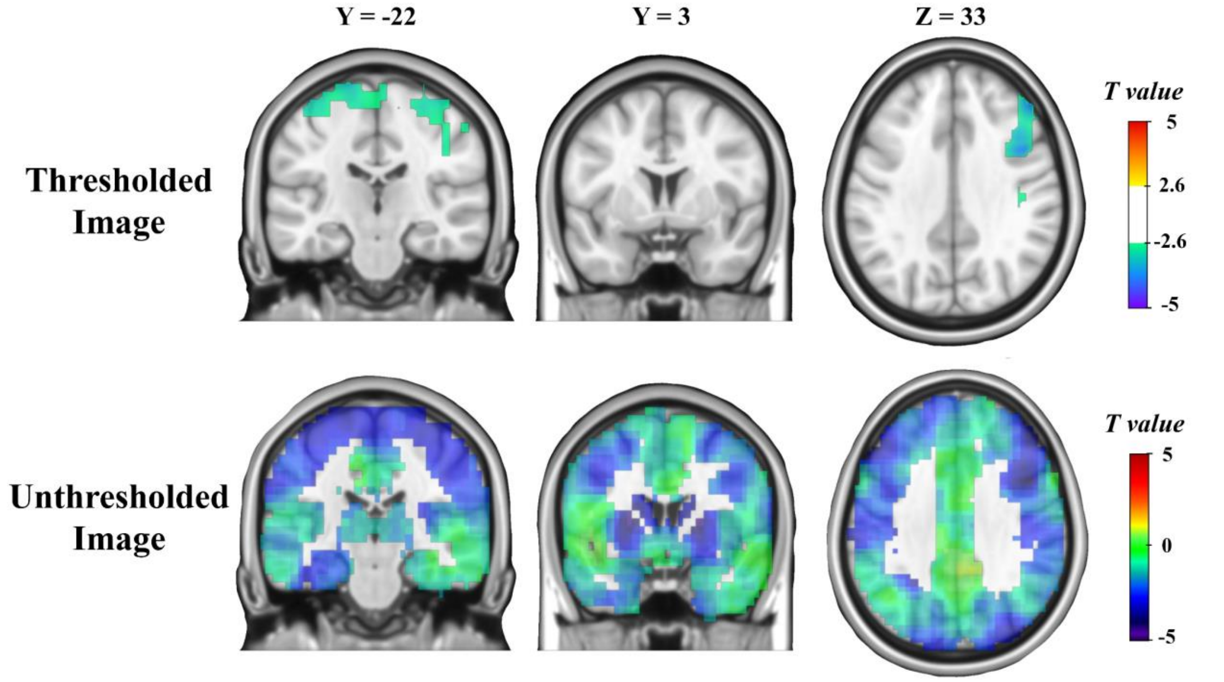


**Supplementary Figure 3.** Thresholded and unthresholded maps of T-values from a regression of eSPPB against brain entropy maps in the cohort of older adults. Cooler colors indicate a negative association between eSPPB and entropy. The regression controlled for the number of volumes removed from scans due to head motion.

Supplementary Figure 4 shows results from a regression of BMI against entropy maps while adjusting for in-scanner motion. Results were very similar to Figure 3 from the main paper.

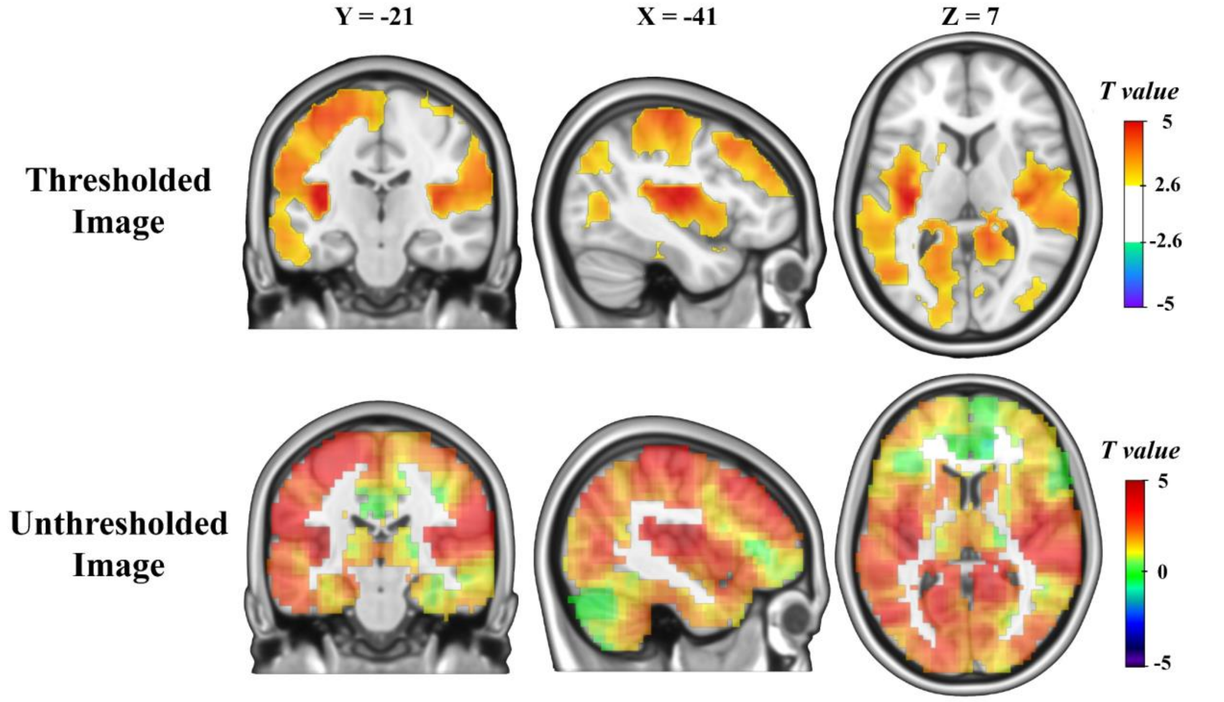


**Supplementary Figure 4.** Thresholded and unthresholded maps of T-values from a regression of BMI against brain entropy maps in the cohort of older adults. Warmer colors indicate a positive association between BMI and entropy while cooler colors indicate a negative relationship. The regression controlled for the number of volumes removed from scans.

Given the strength of the difference between the older and younger cohorts (Figure 1 and Supplementary Figure 2), we evaluated whether age was associated with entropy within the older sample. A regression model was fit in the older adult sample with age as the sole regressor and entropy maps as the outcome. Supplementary Figure 5 shows results from this regression. No clusters remained significant following correction for multiple comparisons – therefore, only unthresholded images are shown. The regions that exhibited the largest, though not statistically significant, associations between age and entropy were located in the dorsal parietal, occipital, and medial prefrontal cortex, as well as in the basal ganglia.

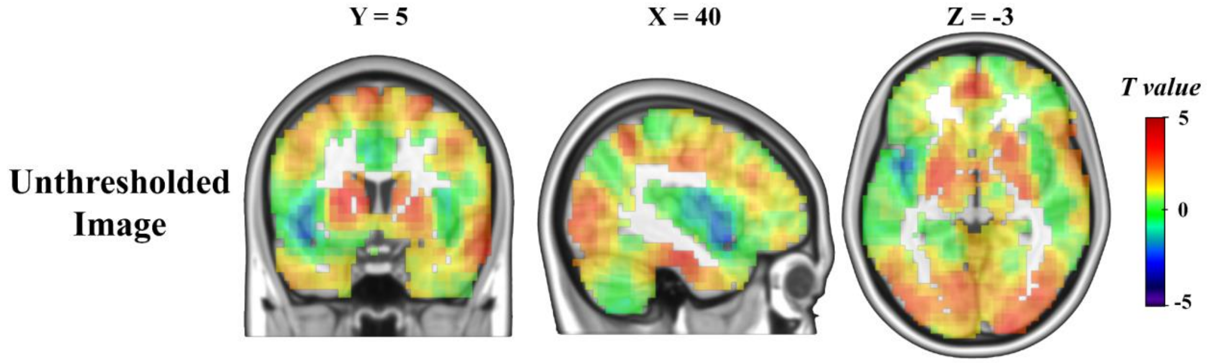


**Supplementary Figure 5.** Unthresholded maps of T-values from a regression of age against brain entropy maps in the cohort of older adults. Warmer colors indicate a positive association between eSPPB and entropy while cooler colors indicate a negative relationship.

In further exploratory analyses, we evaluated several other measures of general health. Results from these exploratory analyses are shown in Supplementary Figures 6-8. Supplementary Figure 6 shows results from a T-test of older participants with and without type 2 diabetes history. Participants with type 2 diabetes had significantly higher entropy in a cluster of voxels spanning from the midcingulate to the left sensorimotor cortex.

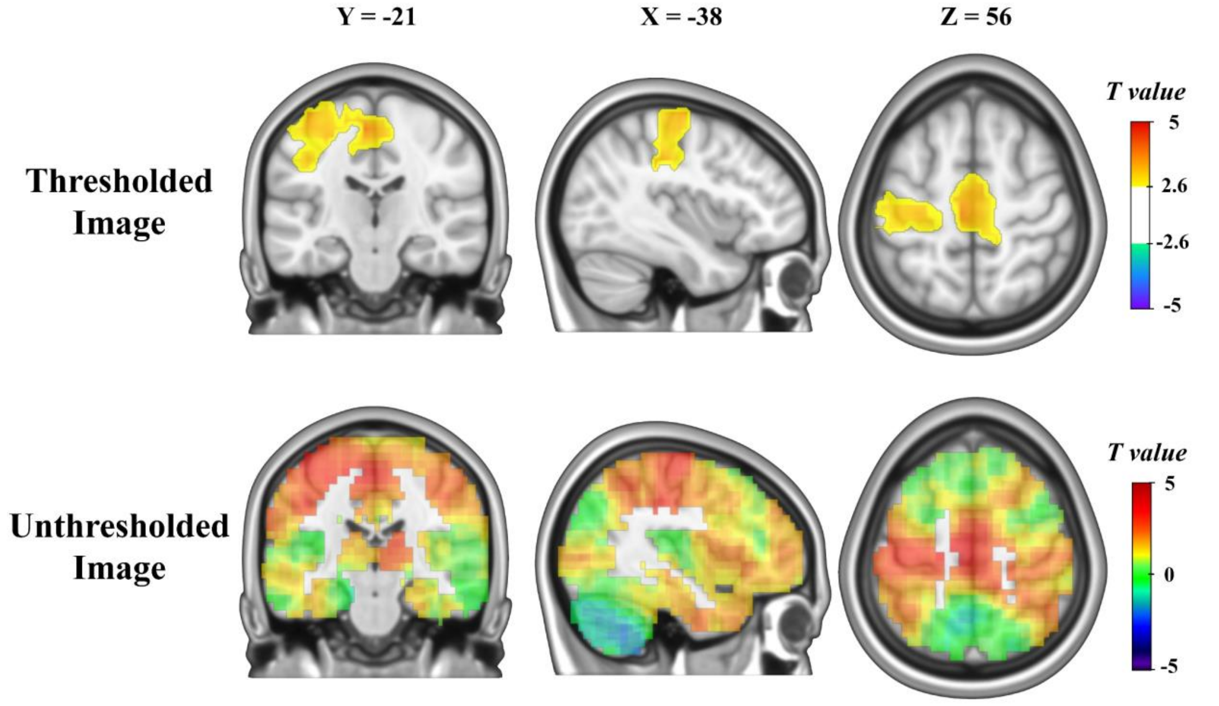


**Supplementary Figure 6.** Thresholded and unthresholded maps of T-values from a two-sample t-test comparing entropy maps of older adults with and without diabetes history. Warmer colors indicate regions with higher entropy in older adults with diabetes history.

Supplementary Figure 7 shows results from a T-test of older participants with and without hypertension. Participants with hypertension had significantly higher entropy in the majority of the occipital lobe.

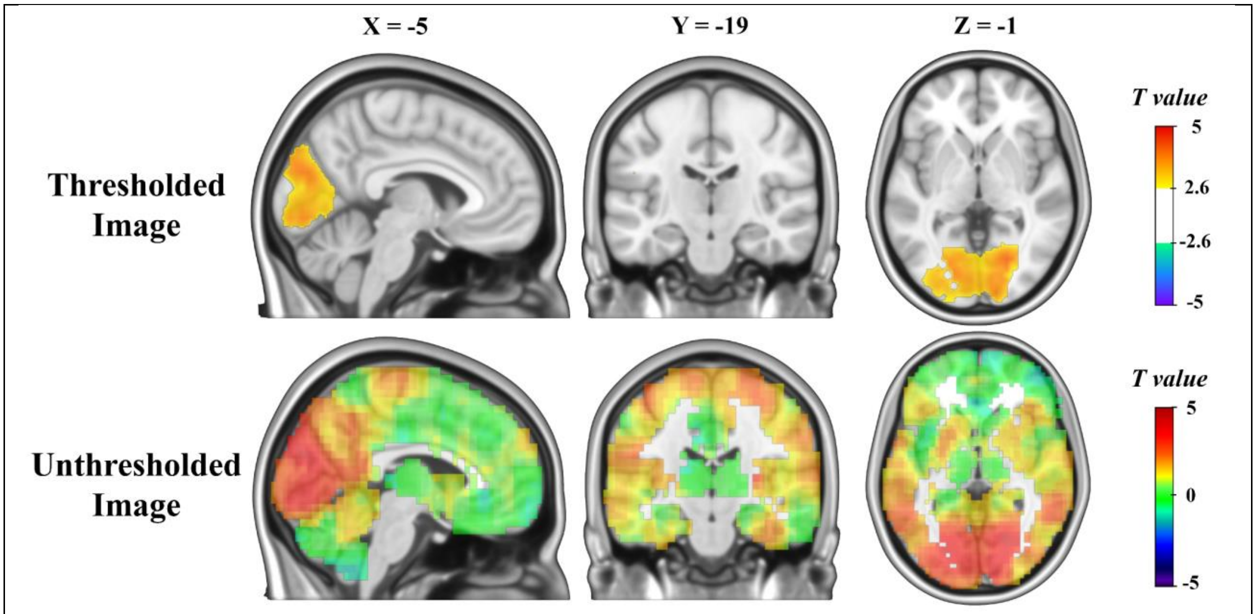


**<u>Supplementary Figure 7.</u>** Thresholded and unthresholded maps of T-values from a two-sample t-test comparing entropy maps of older adults with and without hypertension history. Warmer colors indicate regions with higher entropy in older adults with hypertension.

Supplementary Figure 8 shows results from a two-sample t-test of older participants who did or did not report exercising at moderate intensity for at least 150 minutes per week. Physically active individuals had significantly lower entropy in the basal ganglia and portions of the left and right hippocampus.

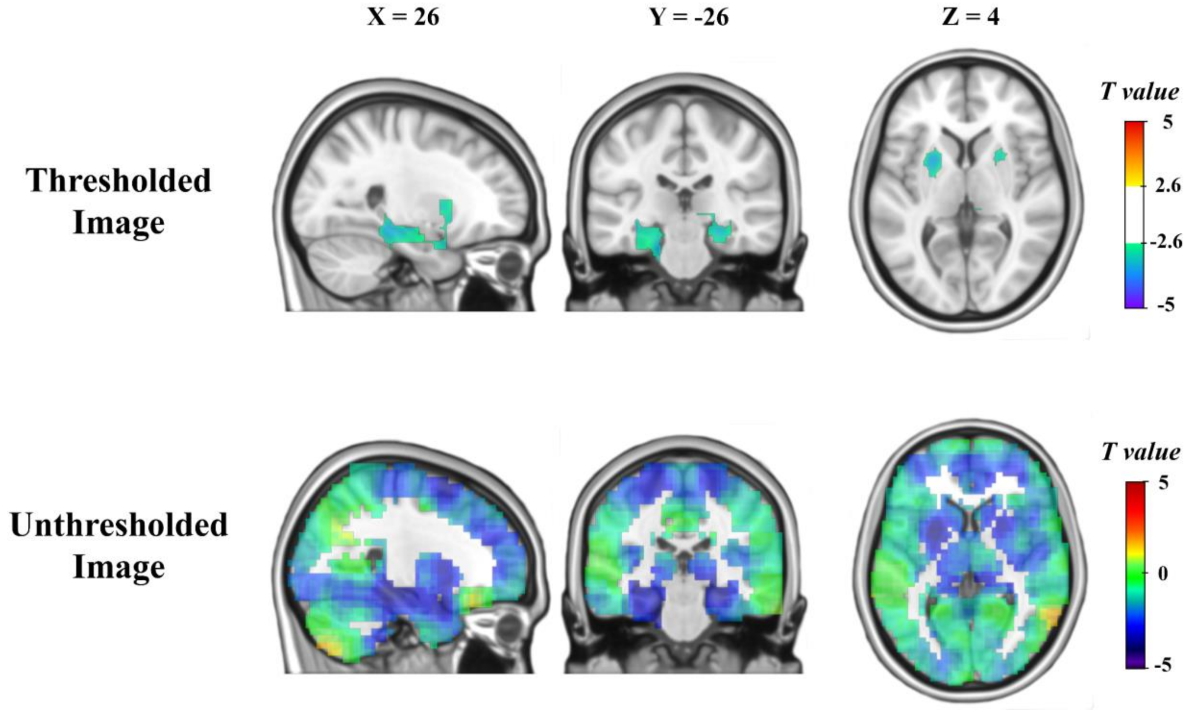


**<u>Supplementary Figure 8.</u>** Thresholded and unthresholded maps of T-values from a two-sample t-test comparing entropy maps of older adults who do meet physical activity guidelines versus older adults who do not. Cooler colors indicate regions with higher entropy in the physically inactive older adults.

Stewart, A. L., Mills, K. M., King, A. C., Haskell, W. L., Gillis, D., & Ritter, P. L. (2001). CHAMPS physical activity questionnaire for older adults: outcomes for interventions. *Med Sci Sports Exerc*, *33*(7), 1126-1141. https://doi.org/10.1097/00005768-200107000-00010
